# Effect of Assembly Method on Nanoparticle Attachment Density, Growth Rate, and Motility of Nanoscale Bacteria Enabled Autonomous Drug Delivery System (NanoBEADS)

**DOI:** 10.1101/867101

**Authors:** Ying Zhan, Austin Fergusson, Lacey R. McNally, Richey M. Davis, Bahareh Behkam

## Abstract

Microbial-mediated drug delivery systems have the potential to significantly enhance the efficacy of nanomedicine for cancer therapy through improved specificity and interstitial transport. The Nanoscale Bacteria-Enabled Autonomous Drug Delivery System (NanoBEADS) is a bacteria-based bio-hybrid drug delivery system designed to carry nanotherapeutics cargo deep into poorly vascularized cancerous tissue. The effect of bacteria-nanoparticle conjugation method and NanoBEADS assembly parameters (*i.e.*, mixing method, volume, and duration) was investigated to maximize particle attachment density. The nanoparticle attachment capacity, viability, growth rate and motility of the original NanoBEADS and an antibody-free variant NanoBEADS were characterized and compared. It is found that the assembly parameters affect the attachment outcome and the binding mechanism impacts the attachment number, the growth rate and motility of NanoBEADS. The NanoBEADS platform provides an opportunity to load nanoparticles with different materials and sizes for applications beyond cancer therapy, such as imaging agents for high-resolution medical imaging.

## Introduction

The advancement of three-dimensional micro/nano fabrication, sensing, and actuation technologies has paved the way for applications of microrobots in medicine. In contrast to the traditional systemic administration of therapeutics, microrobots are capable of targeted drug delivery. Host cells, attenuated pathogens, and GRAS microorganisms are exploited as *living machines* for transport of therapeutic loads. When integrated with synthetic materials, these are called biohybrid microrobots. Stem cells [1], muscle cells [2], immune cells [3, 4], red blood cells [5], bacteria [6, 7] and algae [8], are typical biological materials used in biohybrid microrobots as on-board sensors and actuators. The sensing and actuation are powered by chemical energy. Such biohybrid materials have significant potential for targeted delivery of drugs, genes, mRNA, proteins, imaging contrast agents, radioactive seeds and stem cells to specific locations accessible via the vasculature and even interstitially through self-propulsion. Bacteria possess unique properties of self-propulsion and taxis behaviors (chemotaxis [6, 7], aerotaxis [9], magnetotaxis [10], phototaxis, and pH-taxis [11]), are easy to genetically manipulate to produce attenuated strains with a good safety profile. The advantages of targeted accumulation, deep penetration, and easy genetic modification make bacteria ideal candidates for therapeutic delivery agents.

Bacteria-based bio-hybrid drug delivery systems are comprised of living bacteria for sensing and controlled transport and synthetic micro- or nanoparticles as cargo. An effective bio-hybrid drug delivery system relies on a suitable attachment mechanism and an appropriate distribution of cargo on the cell surface. The methods of particle attachment and the particle attachment density can affect bacterial motility and subsequently the efficacy of the drug delivery agent [21]. The attachment mechanism between bacteria and cargo can also affect the stability of particle attachment. A simple attachment method utilizes electrostatic attraction between the negatively charged outer membrane of the bacteria and the positively charged surfaces of cargo nanoparticles [12–14]. Some other common methods involve hydrophobic interactions [15] and bioaffinity interactions to achieve attachment. Bioaffinity interactions can involve streptavidin-biotin [16, 17], avidin-biotin [5], antibody-antigen [18], or combination of streptavidin-biotin and antibody-antigen [5, 19–21]. Covalent interactions such as the crosslinking of amine group and carboxylate groups can be used [22–24]. Electroporation has also been used to cause drug-containing liposomes to be internalized in bacteria [14, 25].

Previously, we developed a bacteria-based bio-hybrid drug delivery system called Nanoscale Bacteria-Enabled Autonomous Drug Delivery System (NanoBEADS) [21]. NanoBEADS consist of the tumor-targeting *S.* Typhimurium VNP20009 to which nanoparticles comprised of a biodegradable copolymer, poly (lactic-co-glycolic acid) (PLGA), are attached. The particles were first functionalized with streptavidin using 1-ethyl-3-(-3-dimethylaminopropyl) carbodiimide hydrochloride (EDAC) coupling chemistry and the bacteria were coated with a biotinylated antibody. Upon mixing these suspensions, the high affinity between biotin and streptavidin resulted in the attachment of the nanoparticles to the bacteria. The resulting NanoBEADS can autonomously deliver and distribute nanoparticles within the tumor microenvironment *in vitro* and *in vivo* [26]. The nanoparticle quantity per bacterium, viability, and growth rate of the original NanoBEADS was characterized but the effects of variations on the assembly method parameters – specifically mixing volumes during the biotin-streptavidin coupling steps, the method of mechanical mixing during the coupling step, and a different method for attaching biotin to the bacterial surface – have not been investigated until now. Furthermore, the effect of nanoparticle size on the its bacterial attachment density, and NanoBEADS growth rate and motility is largely unexplored.

In this work, we investigate the effects of different attachment methods on NanoBEADS properties (Figure 1). First, we investigated the effect of NanoBEADS assembly parameters (*i.e.*, mixing method, volumes, and times) to maximize particle attachment density. Based on the original NanoBEADS concept, a variant of NanoBEADS was developed: antibody-free NanoBEADS. To produce antibody-free NanoBEADS, bacteria were first mixed with a solution of biotin so that some of the biotin would physically adsorb to the cells. Streptavidin-coated nanoparticles were then attached to the bacteria using the two optimal attachment methods. We characterized and compared the nanoparticle attachment density, viability, growth rate and motility of the variants of NanoBEADS. We found the assembly parameters affect the attachment outcome and the binding mechanism impacts the attachment number, the growth rate and motility of NanoBEADS.

**Figure 1.**
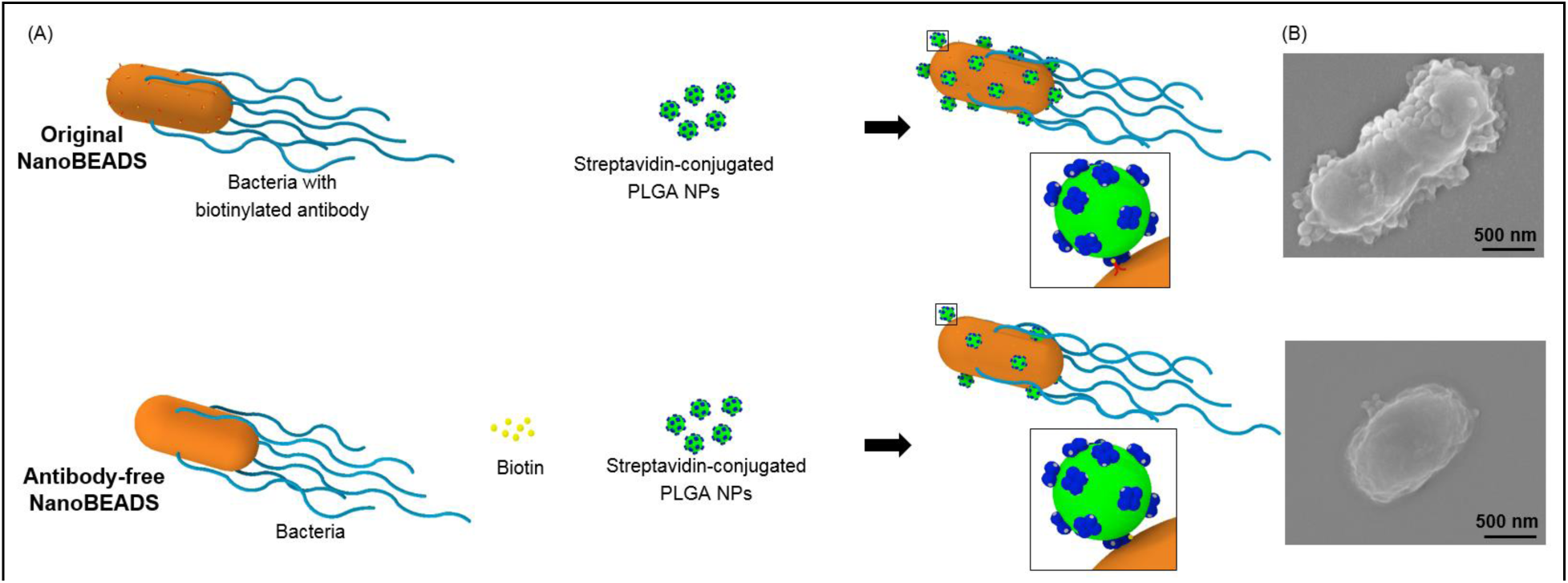
Nanoscale Bacteria-Enabled Drug Delivery System (NanoBEADS). Schematic (A) and representative scanning electron microscopy (SEM) images (B) of the original and variants of NanoBEADS.

## Results and Discussion

### Effect of assembly method on nanoparticle attachment density

The nanoparticle attachment density is important in that it is expected to affect the efficacy of bacteria-based drug delivery systems. We first investigated the effect of agitation method (*i.e.*, the type of mechanical mixer employed when mixing the streptavidin-coated nanoparticles (NPs) and the biotinylated bacterial suspensions), assembly volume (*i.e.*, total suspension volume during mixing), and assembly time, as depicted in Figure 2(A), on nanoparticle attachment outcome of the NanoBEADS. The number of attached nanoparticles on each bacterium was quantified through SEM imaging and reported as the number of attached NPs per projected unit area of cell surface. As shown in Figures 2(B) and (C), assembly volume had no significant effect on the attachment yield of NanoBEADS constructed using a vortex mixer, except for the 60 mins assembly period. For a given assembly period, the nutating mixer and end-over-end mixer resulted in higher attachment numbers compared to the vortex mixer. As the assembly time was increased from 30 mins to 60 mins, the count of NanoBEADS with attachment numbers higher than 10 NPs/µm^2^ cell area showed an increase, especially using the nutating mixer and end-over-end mixer (Figure 2C). Upon increasing the assembly time to 90 mins, more NanoBEADS with attachment numbers lower than 10 NPs/µm^2^ cell area were obtained, resulting in an overall decrease of the average attachment number. Therefore, 60 mins was identified as the optimal assembly time. Altogether, it was established that the nutating mixer used with 800 µL assembly volume for 60 mins (N-800-60) and the end-over-end mixer used with 800 µL assembly volume for 60 mins (E-800-60) are the optimal assembly parameters, which were further examined for repeatability in nanoparticle attachment density outcomes for the original NanoBEADS as well as a variant.

**Figure 2.**
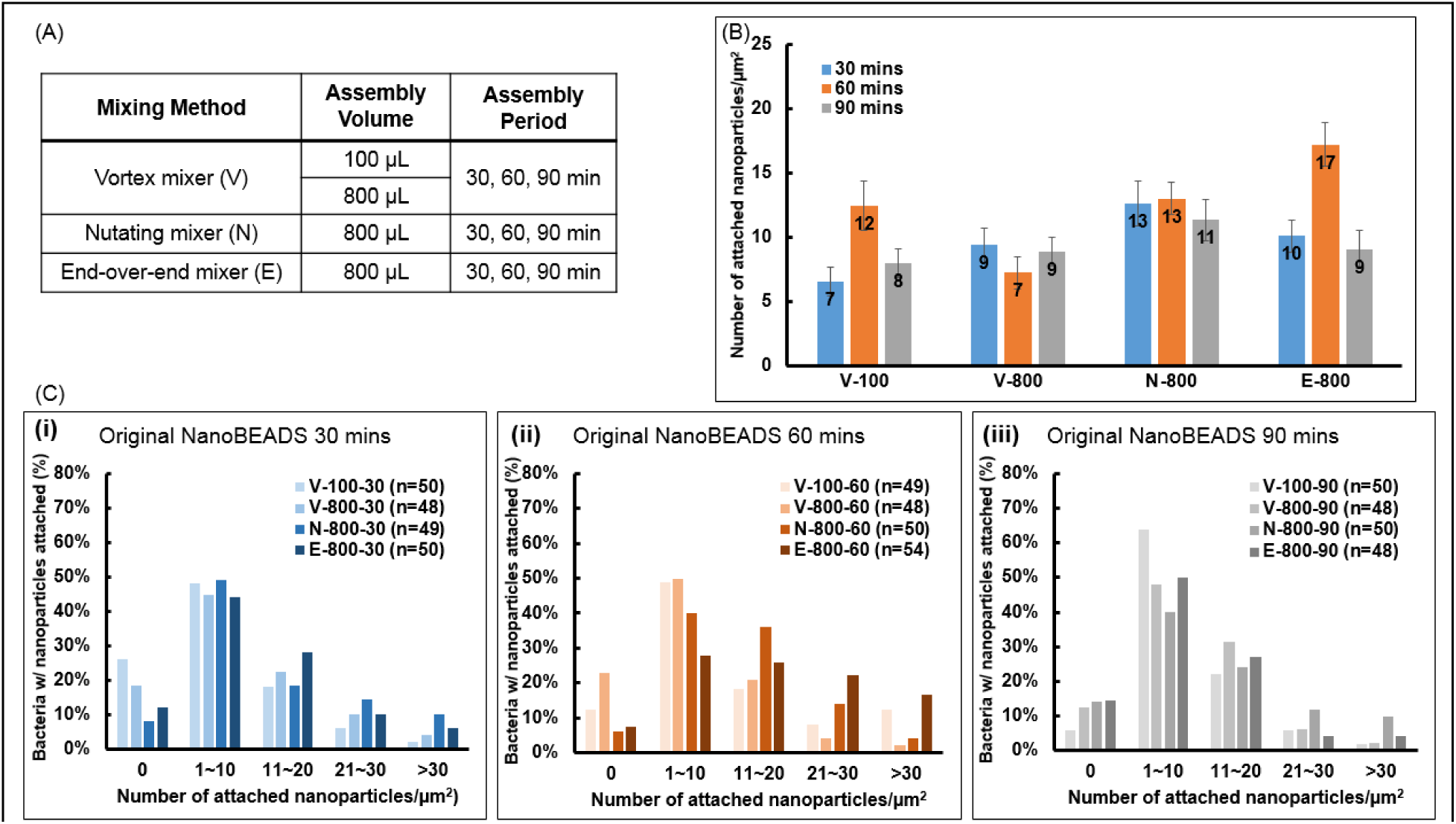
Effect of assembly period, assembly volume, and mixing method on nanoparticle attachment density in the original NanoBEADS. (A) Table of assembly parameter (see Methods for details). (B) The effect of NanoBEADS assembly method and duration on nanoparticle attachment density. (C) Distribution of attached nanoparticles/μm^2^.

A minimum of three independent experiments of original NanoBEADS and antibody-free NanoBEADS were conducted with the assembly methods B-800-60 and E-800-60 (Figure 3). For the original NanoBEADS, both B-800-60 and E-800-60 yielded similar results; however E-800-60 produced more consistent attachment results than B-800-60. Thus, we identify E-800-60 as the optimal set of assembly parameters for producing original NanoBEADS. For antibody-free NanoBEADS, no discernible difference between B-800-60 and E-800-60 outcomes was observed. Comparing the distribution of attached nanoparticles numbers, for the original NanoBEADS, 87% or more bacteria had some particle attachment, while 66% or more of antibody-free NanoBEADS had some particle attachment. It is evident that the original NanoBEADS had a higher number bacteria with greater attachment number than antibody-free NanoBEADS. The viability of the bacteria was not affected by the attached nanoparticles (**Figure S1**).

**Figure 3.**
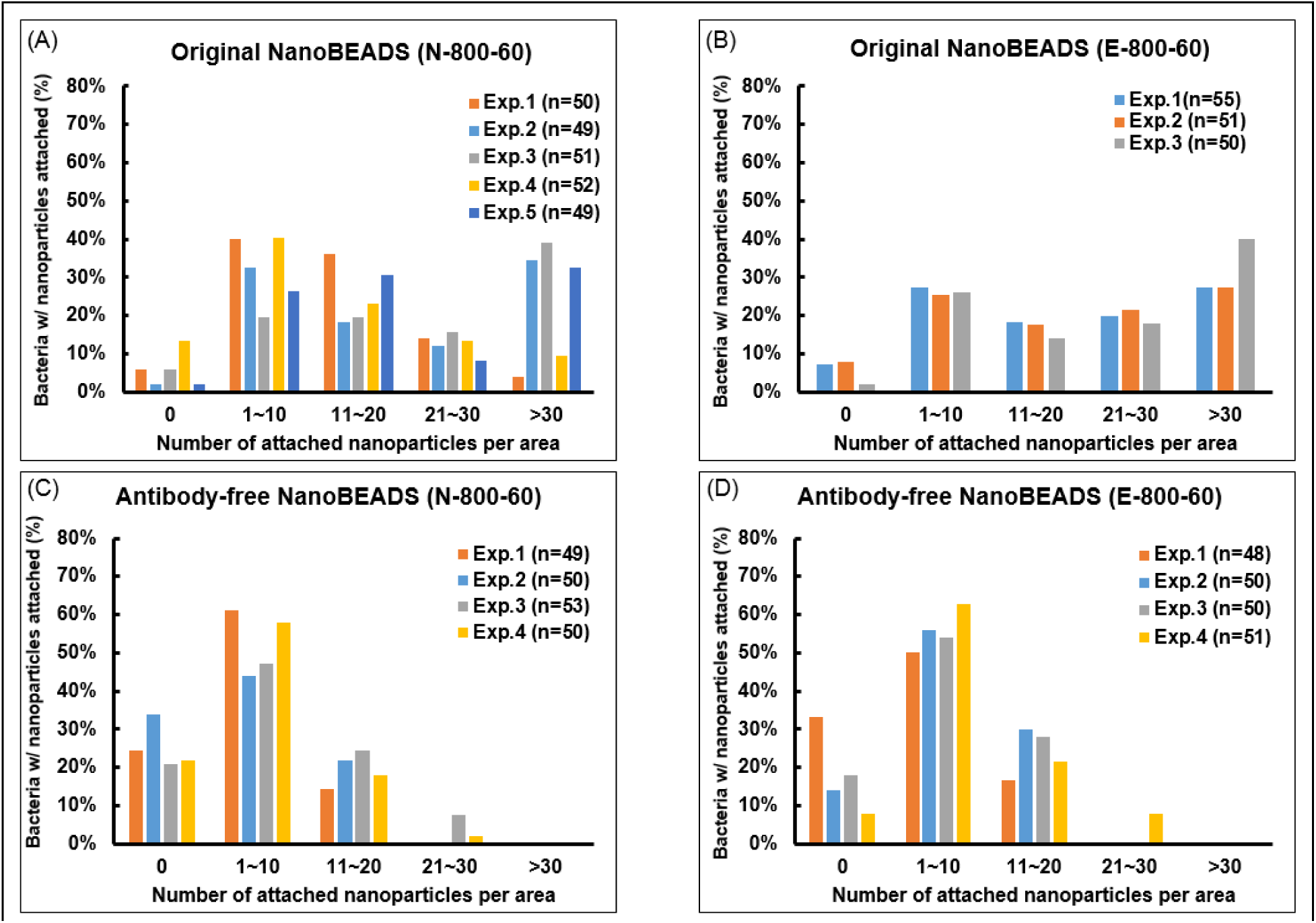
Effect of assembly and agitation methods on nanoparticle attachment density. (A) The repeatability of original NanoBEADS. (B) The comparison of the distribution of attached nanoparticles number of original NanoBEADS. (C) The repeatability of antibody-free NanoBEADS. (D) The comparison of the distribution of attached nanoparticles number of antibody-free NanoBEADS.

The average numbers of nanoparticles attached to bacteria for the original NanoBEADS and antibody-free NanoBEADS are shown in Figure 4(A). There was a 70% decrease in the number of attached particles/μm^2^ upon changing from the original NanoBEADS approach to the antibody-free binding method for both the E-800-60 and B-800-60 cases. In conclusion, the method of attaching biotin to the bacteria affects the NP attachment outcome. When biotin is attached using the biotinylated antibody, more particles per bacteria are attached compared to using physisorbed biotin.

**Figure 4.**
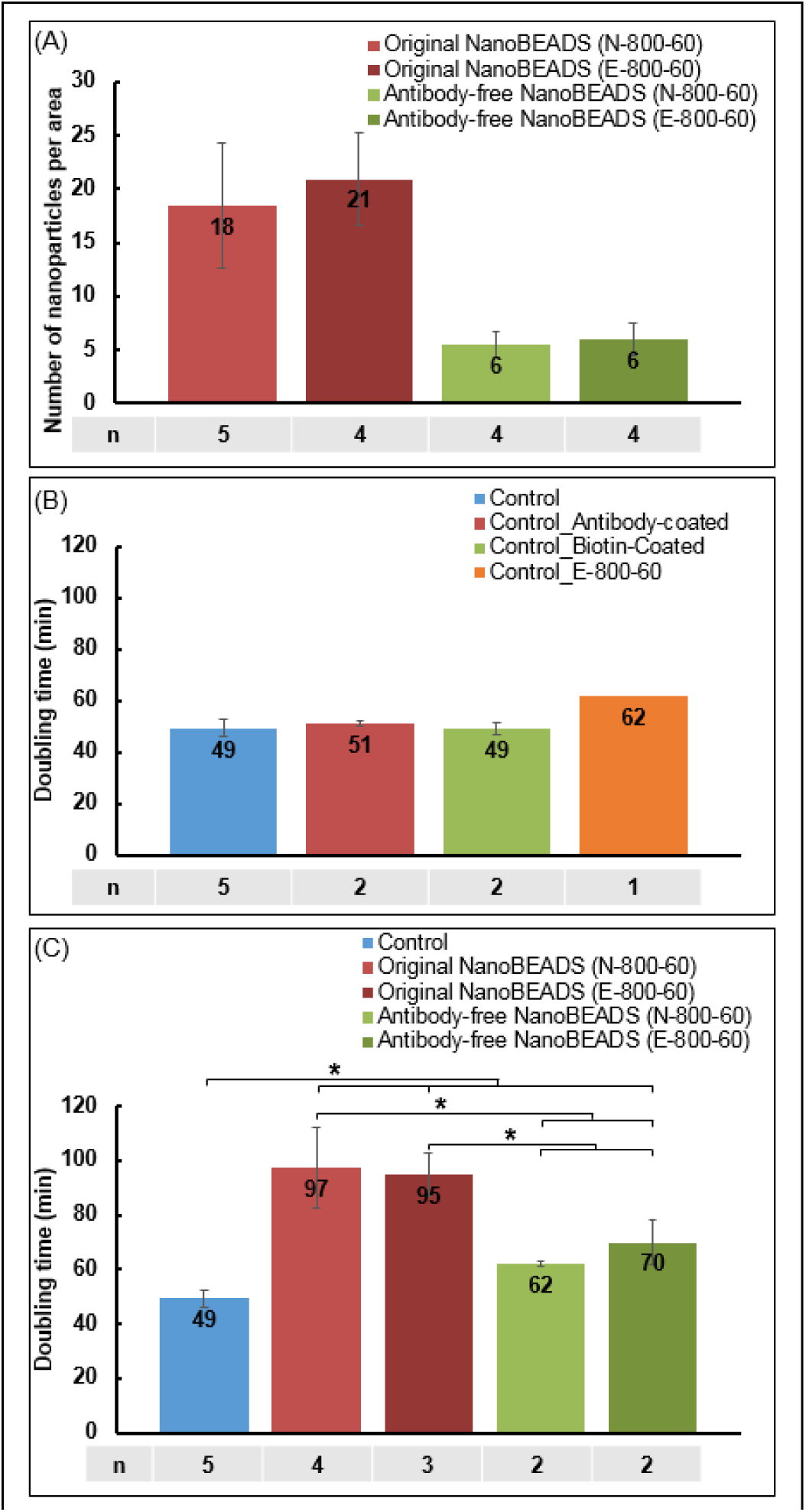
(A) The average number of attached nanoparticles of all variants of NanoBEADS. (B)The comparison of doubling time of the bacteria, antibody-coated bacteria, biotin-coated bacteria and mechanically treated bacteria(E-800-60) with no particles attached. (C)The comparison of doubling time of the variants of NanoBEADS.

### Effect of assembly method on growth rate

To determine the effect of assembly method on the growth of NanoBEADS, we measured the doubling time of bacteria and NanoBEADS. All experiments were performed in tumor culture medium, given the ultimate application of the NanoBEADS platform as a cancer drug delivery system. For a given assembly method, E-800-60, there was no difference of average doubling time between unmodified bacteria, antibody-coated bacteria and biotin-coated bacteria, which means surface modification had no effect on the bacterial growth rate (Figure 4(B)). However, subjecting the bacteria to the mechanical agitation experienced during the E-800-60 assembly process did increase the average doubling time for the unmodified bacteria from 49 mins to 62 mins which is reasonable given the additional stress imposed on the organisms. Next, we measured the NanoBEADS growth rate. The average doubling times of the original NanoBEADS using methods B-800-60 and E-800-60 were 97 mins and 95 mins, respectively, as compared to 49 mins for the unmodified bacteria (Figure 4(C)). The average doubling time of antibody-free NanoBEADS (B-800-60) and antibody-free NanoBEADS (E-800-60) were 62 mins and 70 mins. The bacteria doubling time also increased consistently with the particle attachment number as seen in Figures **4(A-C)** which is reasonable since the particle attachment imposes stresses on the bacteria. The particular mixing method (*i.e.*, nutating mixer vs. end-over-end mixer) had no discernible effect on the growth rate for both the original NanoBEADS and the antibody-free case as seen in Figure 4(C).

### Effect of assembly method on NanoBEADS motility

We assessed the effect of assembly method on motility by measuring the swimming speeds of bacteria, the original NanoBEADS, and the antibody-free NanoBEADS. The average speeds of bacteria, mechanically treated bacteria (B-800-60 and E-800-60 without particles or antibody coating), and NanoBEADS constructed using B-800-60 and E-800-60 assembly parameters are shown in Figure 5(A-B). Mechanical agitation decreased the average speed of bacteria from 15.3±3.2 µm/s to 11.3±4.0 µm/s (B-800-60) and 10.9±3.9 µm/s (E-800-60), respectively (Figure 5(A)). This is reasonable since agitation could damage bacteria flagella on the bacteria as well as perturb them. There was no significant difference on speed between using the nutating mixer and the end-over-end mixer. The average speed of original NanoBEADS was similar to that of the antibody-free NanoBEADS, despite the significant difference in the number of attached nanoparticles (Figure 5(B)).

**Figure 5.**
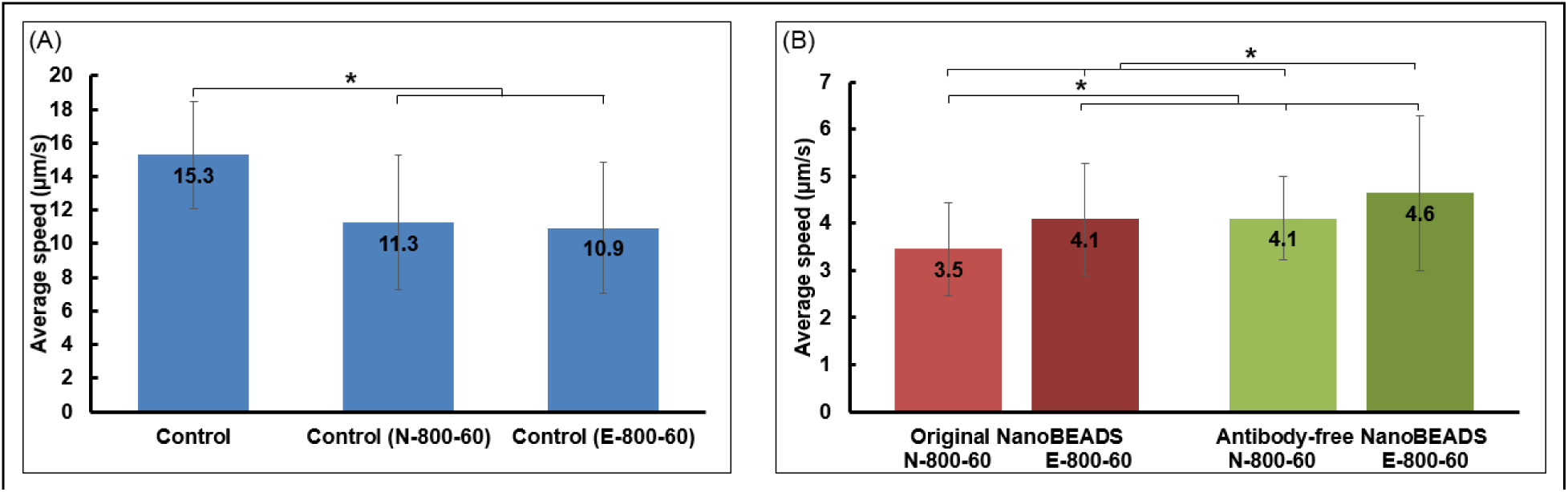
(A)The average speed of bacteria at 2 hrs incubation time as a function of mixing conditions without particles and antibody coating. (B) The average speed of NanoBEADS at 2 hrs incubation time.

Upon characterizing and comparing the nanoparticle attachment capacity, motility and growth rate of the variants of NanoBEADS, we demonstrate that other nanoparticles (i.e., silica and gold nanoparticles) can be attached to bacteria at high attachment capacity using the same procedure. For Silica NanoBEADS and Au NanoBEADS, silica nanoparticles and gold nanoparticles were used instead of PLGA nanoparticles as that in original NanoBEADS (**Figure S2**).

## Conclusions

The NanoBEADS is a bacteria-based bio-hybrid drug delivery system designed to enhance the efficacy of bacteria to deliver nanomedicine. The assembly parameters and binding mechanisms affect the property of system (i.e., attachment outcome, growth rate and swimming speed of bacteria). For the original NanoBEADS, using the end-over-end mixer with a total mixing volume of 800 µL for 60 mins assembly period (E-800-60) was optimal for achieving maximum attachment outcome without affecting the viability of the bacteria. When streptavidin-functionalized nanoparticles were attached to the bacteria using biotin that was physisorbed to the bacteria, the attachment number decreased by more than 60% compared to the original NanoBEADS case. We also showed that particle attachment reduces the growth rate and the motility of bacteria. Furthermore, we demonstrated that other nanoparticles (i.e., silica and gold nanoparticles), instead of PLGA nanoparticles as that in original NanoBEADS, can be attached to bacteria with promising attachment outcomes (**Figure S2**). The NanoBEADS platform provides the possibility to load other nanoparticles with different materials and sizes for various applications in cancer therapy such as imaging agents for high-resolution medical imaging.

## Acknowledgements

This project was partially supported by the National Science Foundation (CAREER award, CBET-1454226) and the Institute for Critical Technology and Applied Science (ICTAS) at Virginia Tech.

## Methods

### Bacteria preparation

*Salmonella* Typhimurium VNP20009 *cheY*^*+*^ [27] from a single colony was cultured overnight in MSB medium (10 g/L tryptone, and 5 g/L yeast extract, 2 mM MgSO_4_, 2mM CaCl_2_, pH 7.0) at 37°C with shaking at 100 rpm (Excella E24 Incubator Shaker Series, New Brunswick Scientific). The overnight bacterial culture was diluted in MSBto 1% and incubated at 37°C with shaking at 100 rpm until an optical density at 600 nm (OD_600_) of 1.0 was reached. The bacterial culture was centrifuged at 1,700 g for 5 minutes twice and re-suspended in motility media (6.4 mM K_2_HPO_4_, 3.5 mM KH_2_PO_4_, 1 μM L-methionine, 10 mM Sodium DL-lactate, 2 mM MgSO_4_, 2 mM CaCl_2_, pH 7.0) to an OD_600_ of 1.0.

### PLGA Nanoparticle Synthesis

100 mg of Pluronic F127 and 20 mL of deionized water were added to a glass vial. The vial was placed in a water bath sonicator (Branson 2510 Ultrasonic Cleaner, 100 W) for 30 minutes to dissolve the Pluronic then a magnetic stir bar was added, and the solution was stirred at 600 rpm. Acid-terminated PLGA (Mw: 25,000 g mol^−1^, 50:50 lactic acid:glycolic acid, acid end‐capped, Akina Inc. PolySciTech, West Lafayette, IN) was dissolved with dimethylformamide (DMF, Sigma‐Aldrich, St. Louis, MO) to a final concentration of 22.22 mg/mL; the solution was sonicated for 30 minutes to ensure molecular dissolution. While the PLGA solution was sonicating, 6,13‐bis(triisopropylsilylethynyl) pentacene (TIPS, Sigma‐Aldrich, St. Louis, MO) was dissolved in tetrahydrofuran (THF, anhydrous and uninhibited, >99.9%, Sigma‐Aldrich, St. Louis, MO) to a final concentration of 3.05 mg/mL. After the PLGA solution was fully dissolved, TIPS solution was added to the PLGA solution to achieve a 1:10 THF:DMF volume ratio. The mixture was vortexed for ~5 seconds before it was loaded into a 5 mL glass syringe with a 21-gauge needle attached. Care was taken to remove all macroscopic air bubbles from the syringe. The TIPS:PLGA mixture (1 mL) was added dropwise (0.5 mL/min, NE-1000, New Era Pump Systems Inc.) to the stirring (600 rpm) Pluronic F127 solution. The resulting nanoparticle suspension was allowed to stir for 5 hours at 600 rpm. The suspension was protected from light to prevent degradation of the fluorophore. After 5 hours, the suspension was centrifuged at 22,789 xg for 30 minutes at 4ºC (Sorvall Legend X1R, Thermofisher Scientific). The supernatant was discarded. The pellet was resuspended in 20 mL of 1x PBS by vortex mixing for 2 minutes and sonicating the suspension for 30 minutes. The final dispersed suspension was passed through a nitrocellulose syringe filter (0.45 µm pore size) to remove any remaining aggregates. The filtrate was stored in a foil-wrapped vial at room temperature.

### Streptavidin Functionalization of PLGA Nanoparticles

Microcentrifuge tubes were filled with 700 µL of PLGA NP suspension. The tubes were centrifuged at 16,060 xg for 10 minutes at room temperature (accuSpin Micro, Fisher Scientific). The pellets were resuspended in 800 µL of EDAC coupling solution (20 mg/mL EDAC, 5 µg/mL streptavidin-Cy3, pH 5.2 50 mM MES buffer). The streptavidin coupling reaction took place on a vortex mixer (500 rpm, Fisher Digital Vortex 120V, Fisher Scientific) for 3 hours. Following streptavidin coupling, the microcentrifuge tubes were centrifuged at 16,060 xg for 10 minutes at room temperature. The pellets were resuspended in 100 µL of motility media. Nanoparticle tracking analysis (NTA) measurements were used to determine the nanoparticle concentration in the final suspension prior to incubating the nanoparticles with bacteria to form NanoBEADS.

### Dynamic Light Scattering (DLS) Measurements

DLS measurements were performed using a Zetasizer Nano ZS (Malvern Instruments) operating with Zetasizer Software v7.12. Disposable polystyrene cuvettes were filled with 1 mL of the final aqueous nanoparticle suspensions. Measurements were performed at 25 °C.

### Zeta Potential Measurements

The final aqueous nanoparticle suspensions were loaded into disposable polystyrene capillary cells. Zeta potential measurements were performed using a Zetasizer Nano ZS at 25 °C.

### Nanoparticle Tracking Analysis (NTA)

Dilutions (10x and 100x) of the functionalized nanoparticle suspensions were analyzed using nanoparticle tracking analysis using a NanoSight NS500 (Malvern Instruments) operating with NanoSight NTA v3.4. All measurements were performed at 25ºC. Five 1-minute videos were taken for each sample, and the nanoparticle scattering cones were tracked to determine the number concentrations of the nanoparticle suspensions.

### PLGA NanoBEADS assembly

To prepare the original NanoBEADS, the prepared bacterial suspension was incubated with 10 µg/mL biotinylated *Salmonella* polyclonal antibody (Thermo Scientific, Waltham, MA, USA) on a vortex mixer at 500 rpm at room temperature for 1 hr. The antibody-coated bacteria suspension was then centrifuged at 1,700 g for 5 minutes to remove free antibody and suspended in motility medium to an OD_600_ of 2.0 (Cary 60 UV-Vis, Agilent Technologies). The suspension of biotinylated antibody-coated bacteria was mixed with the streptavidin-coated nanoparticle at a bacteria to particles ratio of 1:100 with volume 100 µL or 800 µL (as described in Figure 2(A)) and incubated on vortex mixer at 500 rpm, nutating mixer (Nutating Mixer 88861043, Fisher Scientific) at 100 rpm or end-over-end mixer (Multi-Purpose Tube Rotator 88861049, Fisher Scientific) at 15 rpm for 30 minutes, 60 minutes or 90 minutes to facilitate the assembly of nanoparticles onto the bacteria. For antibody-free NanoBEADS, the bacterial suspension was incubated with 1 mg/mL biotin (Fisher BioReagents, Fair Lawn, NJ) on a vortex mixer at 500 rpm at room temperature for 30 minutes to coat bacteria with biotin. Subsequently, the suspension was centrifuged at 1,700 g for 5 minutes to remove free biotin and re-suspended in motility medium to an OD_600_ of 2.0. The suspension of bacteria coated with biotin was mixed with the streptavidin-coated nanoparticles at a bacterium to particles ratio of 1:100 with volume 800 µL and incubated on nutating mixer at 100 rpm or end over end mixer at 15 rpm for 60 minutes. After the assembly process, the suspension of NanoBEADS was transferred to a centrifugal filter of 0.8 µm pore size high-flux polyethersulphone membrane (Sartorius Vivaclear, Elk Grove, IL) and centrifuged at 1,700 g for 30 seconds to remove free nanoparticles. The NanoBEADS were suspended in motility medium to an OD_600_ of 1.0.

### NanoBEADS samples preparation for field emission scanning electron microscope (FE-SEM)

To quantify the number of nanoparticles attached on the outer membrane of bacteria, SEM images of NanoBEADS were taken by Leo Zeiss field emission scanning electron microscope (FESEM). Ten µL NanoBEADS solution were deposited on 0.005% (w/v) poly-L-lysine (PLL) treated glass slides and incubated at room temperature for 5 minutes to allow for attachment. Afterwards, the slide was rinsed in DI water to remove the loosely attached nanoparticles and NanoBEADS. Then, the slide was covered with 4% glutaraldehyde for 2 hours at 4°C to fix the attached NanoBEADS. The slide was soaked in 0.1 M Phosphate-buffered saline (PBS) for 20 minutes twice. The same soaking process was repeated with deionized water. After air-drying overnight, the slide was sputter-coated with 7 nm Pt/Pd prior to imaging (Leica ACE600 sputter). High-resolution images were obtained utilizing a LEO (Zeiss) 1550 FESEM at an accelerating voltage of 5 kV, and working distances of <8.6 mm. To determine the average number of attached nanoparticles to each NanoBEADS, the particle numbers on ~ 50 bacteria were counted for each case.

### Viability assay

The filtered NanoBEADS were diluted in motility medium to an OD_600_ of 0.05.To this diluted NanoBEADS solution, 1.5 µL aliquots of the SYTO 9 nucleic acid stain, and propidium iodide (Thermo Fisher, Eugene, Oregon, USA) were added and followed by incubation in the dark at room temperature for 15 minutes. The fluorescent microscopy images of NanoBEADS were taken using a Zeiss AxioObserver Z1 inverted microscope equipped with an AxioCam mRM camera at 40× objective. Live cells with an intact membrane stained green, where cells with a damaged membrane or dead cells stained red.

### Growth rate measurement

The NanoBEADS suspensions were diluted in 3 mL of McCoy’s 5A medium supplemented with 10% FBS to a final OD_600_ of 0.001. The diluted NanoBEADS suspension was incubated at 37°C with shaking at 100 rpm for 10 hours. A 100 µL sample was taken every hour, diluted and plated on three 1.5% LB agar plates. Following an overnight incubation at 37° C, the bacteria colonies on the agar plates were counted to determine the NanoBEADS growth rate.

### Swimming speed measurement

The bacterial and NanoBEADS suspensions were diluted in 3 mL of McCoy’s 5A medium supplemented with 10% FBS to a final OD_600_ of 0.001. The diluted suspensions were incubated at 37°C with shaking at 100 rpm for 2 hours. A 10 µL sample was taken, diluted and placed on the glass cover slip. The videos of bacterial movement were taken with a Zeiss AxioObserver Z1 inverted microscope equipped with an AxioCam Hsm camera and 63x oil immersion objective. The videos were analyzed in ImageJ using the MTrackJ plug-in tool. The average swimming speed was calculated by averaging the instantaneous speeds which are moving distances in each time unit divided by the time unit. For each experiment, about 50 bacteria or NanoBEADS were tracked in the 15 s videos with frame rate of 32.8 fps.

