## Supplemental tables and figures for "Effect of Assembly Method on Nanoparticle Attachment Density, Growth Rate, and Motility of Nanoscale Bacteria Enabled Autonomous Drug Delivery System (NanoBEADS)"

**Table of Contents**

Table S1. **PLGA Nanoparticle Characterization Data including Dynamic Light Scattering and Zeta Potential Results**
Table S2. **Silica Nanoparticle Characterization Data including Dynamic Light Scattering and Zeta Potential Results**
Figure S1. **Comparison of Viability of PLGA NanoBEADS**
Figure S2. **The Variants of Original NanoBEADS**
Figure S3. **Dynamic Light Scattering Measurements of Nanoparticle Suspensions Pre- and Post-Functionalization**

**Chitosan Conjugation of Silica Nanoparticles**

Silica nanoparticles suspended in deionized water (20 mL), with an average diameter of 42 ± 4 nm as measured by dynamic light scattering, were combined with ethanol (5 mL), 10.223 mg/mL chitosan in 5% v/v aqueous acetic acid (20 mL), (3-aminopropyl)trimethoxysilane (100 µL) (APTMS, Sigma-Aldrich, St. Louis, MO), and (3-glycidyloxypropyl)trimethoxysilane (100 µL) (GPTMS, Sigma-Aldrich, St. Louis, MO). The mixture was placed on vortex mixer (2500 rpm) for 90 minutes. Following chitosan conjugation, the suspension was centrifuged at 19,837 ×g for 30 minutes. The supernatant was discarded, and the pellet was resuspended in 5 mL of deionized water with 2 minutes of vortex mixing and 60 minutes of sonication (40 kHz). The Chit-Silica nanoparticles had an average diameter of 300 ± 182 nm as measured by dynamic light scattering.

**Streptavidin Functionalization of Chitosan-Coated Silica (Chit-Silica) Nanoparticles**

Microcentrifuge tubes were filled with 700 µL of Chit-Silica NP suspension. The tubes were centrifuged at 16,060 ×g for 30 minutes. The pellets were resuspended in 800 µL of EDAC coupling solution (20 mg/mL EDAC, 5 µg/mL streptavidin-Cy3, pH 5.2 50 mM MES buffer). The streptavidin coupling reaction took place on a vortex mixer (500 rpm) for 3 hours. Following streptavidin coupling, the microcentrifuge tubes were centrifuged at 16,060 ×g for 30 minutes. The pellets were resuspended in 100 µL of motility medium. The Strept-Chit-Silica nanoparticles had an average diameter of 188 ± 32 nm as measured by dynamic light scattering. Nanoparticle tracking analysis (NTA) measurements were used to determine the nanoparticle concentration in the final suspension prior to incubating the nanoparticles with bacteria to form NanoBEADS.

**N-Hydroxysulfosuccinimide (Sulfo-NHS) Functionalization of PLGA Nanoparticles**

A number of microcentrifuge tubes were filled with 700 µL of PLGA NP suspension. The tubes were centrifuged at 16,060 xg for 10 minutes, and the pellets were resuspended in 800 µL of Sulfo-NHS coupling solution (20 mg/mL EDAC, ~22.65 mg/mL Sulfo-NHS, pH 5.2 50 mM MES buffer). Sulfo-NHS functionalization took place on a vortex mixer (500 rpm) for 30 minutes. Following functionalization, the PLGA NP suspension was centrifuged at 16,060 xg for 10 minutes, and the pellet was resuspended in 100 µL of motility media. The Sulfo-NHS-PLGA nanoparticles had an average diameter of 314 ± 150 nm as measured by dynamic light scattering. Nanoparticle tracking analysis (NTA) measurements were used to determine the nanoparticle concentration in the final suspension prior to incubating the nanoparticles with bacteria.

**Silica NanoBEADS**

The original protocol with optimal assembly parameters (E-800-60) was used to conjugate silica nanoparticles on the surface of *S*. Typhimurium VNP20009 bacteria. During the assembly, the bacteria to nanoparticles ratio was kept 1:100. The schematic and representative SEM images are shown in **Figure S2**.

**Au NanoBEADS**

For Au NanoBEADS, streptavidin-coated gold nanoparticles were used instead of PLGA nanoparticles as that in original NanoBEADS with the optimal assembly parameters (E-800-60). During the assembly, the bacteria to particles ratio was kept 1:100. The average diameter of gold nanoparticles is 42 ± 5 nm as measured by dynamic light scattering. The schematic and representative SEM images are shown in Figure S2.

**Table S1. PLGA Nanoparticle Characterization Data including Dynamic Light Scattering and Zeta Potential Results**

| Uncoupled PLGA Nanoparticles (PLGA NP) | | | |
|---|---|---|---|
| Batch | Size (nm) | Polydispersity Index | Zeta Potential (mV) |
| 1 | 148 | 0.09 | -52.7 |
| 2 | 155 | 0.11 | -48.3 |
| 3 | 184 | 0.12 | -43.7 |
| 4 | 166 | 0.09 | -48.8 |
| 5 | 155 | 0.10 | -51.3 |
| 6 | 148 | 0.10 | -59.7 |
| Streptavidin-PLGA Nanoparticles (Strept-PLGA NP) | | | |
| Batch | Size (nm) | Polydispersity Index | Zeta Potential (mV) |
| 1 | 158 | 0.10 | 23.6 |
| 2 | 156 | 0.07 | 24.3 |
| 3 | 173 | 0.12 | 28.5 |
| 4 | 178 | 0.09 | 28.7 |
| Sulfo-NHS-PLGA Nanoparticles (Sulfo-NHS-PLGA NP) | | | |
| Batch | Size (nm) | Polydispersity Index | Zeta Potential (mV) |
| 5 | 420 | 0.62 | -2.5 |
| 6 | 208 | 0.23 | -20.7 |

**Table S2. Silica Nanoparticle Characterization Data including Dynamic Light Scattering and Zeta Potential Results**

| Silica Core Nanoparticles (Silica NP) | | | |
|---|---|---|---|
| Batch | Size (nm) | Polydispersity Index | Zeta Potential (mV) |
| 1 | 38 | 0.22 | -38.0 |
| 2 | 47 | 0.23 | -36.5 |
| 3 | 41 | 0.35 | -28.8 |
| Chitosan-Coated Silica Nanoparticles (Chit-Silica NP) | | | |
| Batch | Size (nm) | Polydisperisty Index | Zeta Potential (mV) |
| 1 | 509 | 0.58 | 54.7 |
| 2 | 218 | 0.39 | 45.2 |
| 3 | 173 | 0.58 | 49.0 |
| Streptavidin-Functionalized Chitosan-Coated Silica Nanoparticles (Strept-Chit-Silica NP) | | | |
| Batch | Size (nm) | Polydispersity Index | Zeta Potential (mV) |
| 1 | 220 | 0.35 | 24.0 |
| 2 | 157 | 0.42 | 18.7 |
| 3 | 187 | 0.50 | 17.0 |

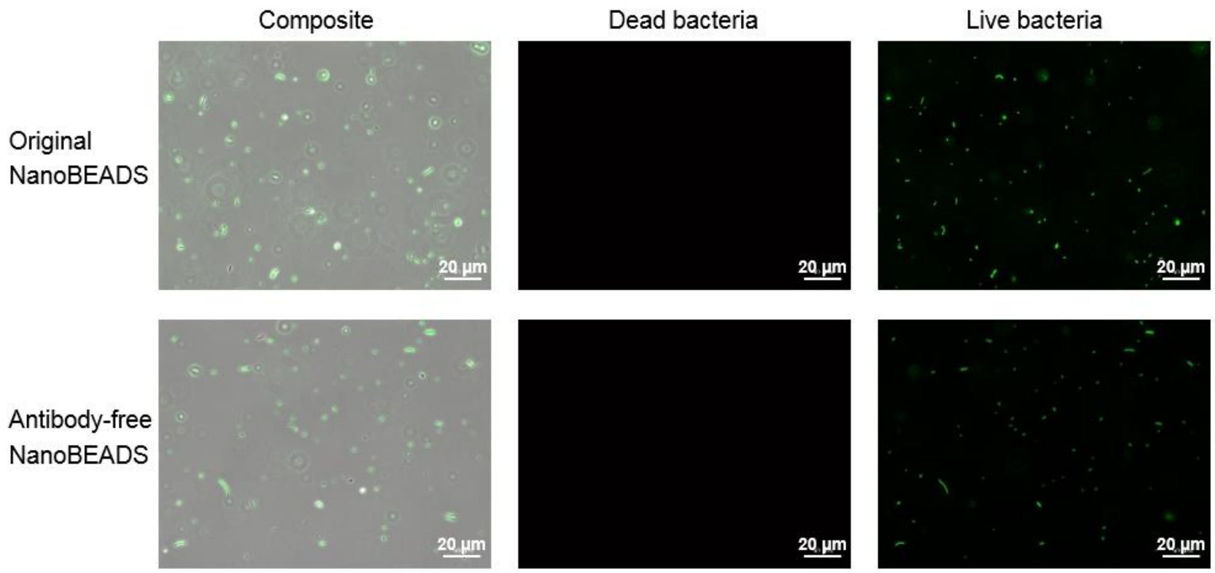


**Figure S1. Comparison of Viability of PLGA NanoBEADS.** A composite bright-field and fluorescence microscopy image of NanoBEADS is shown in the left. The fluorescence microscopy images show dead bacteria (red, middle) and live bacteria (green, right). Nanoparticle conjugation does not affect bacteria viability.

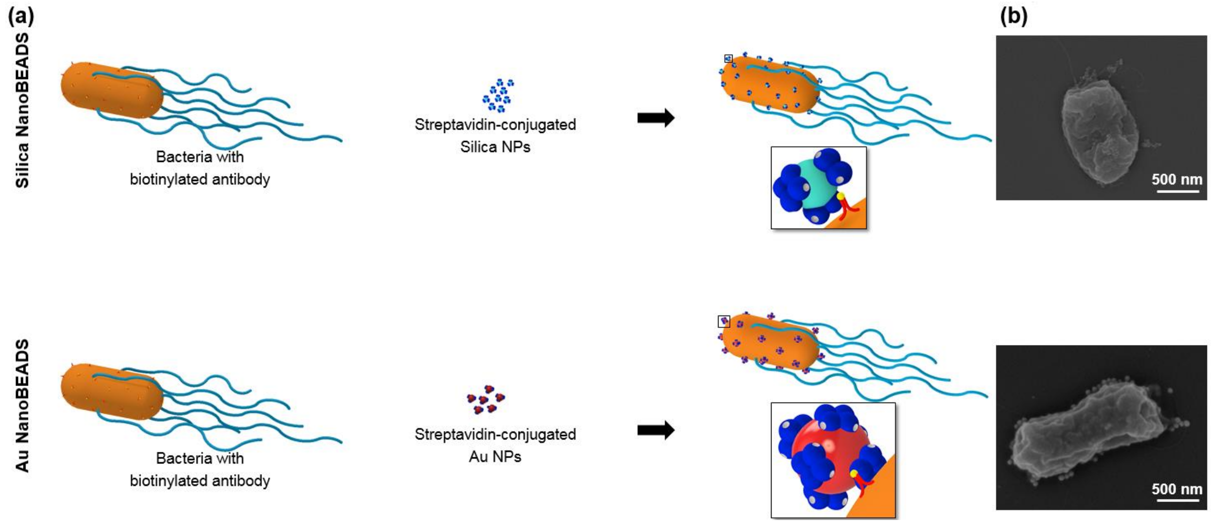


**Figure S2. The Variants of Original NanoBEADS. (a)** Schematic of silica NanoBEADS and Au NanoBEADS. **(b)** Scanning electron microscopy (SEM) images of silica NanoBEADS and Au NanoBEADS. Nanoparticles are false colored for enhanced visibility.

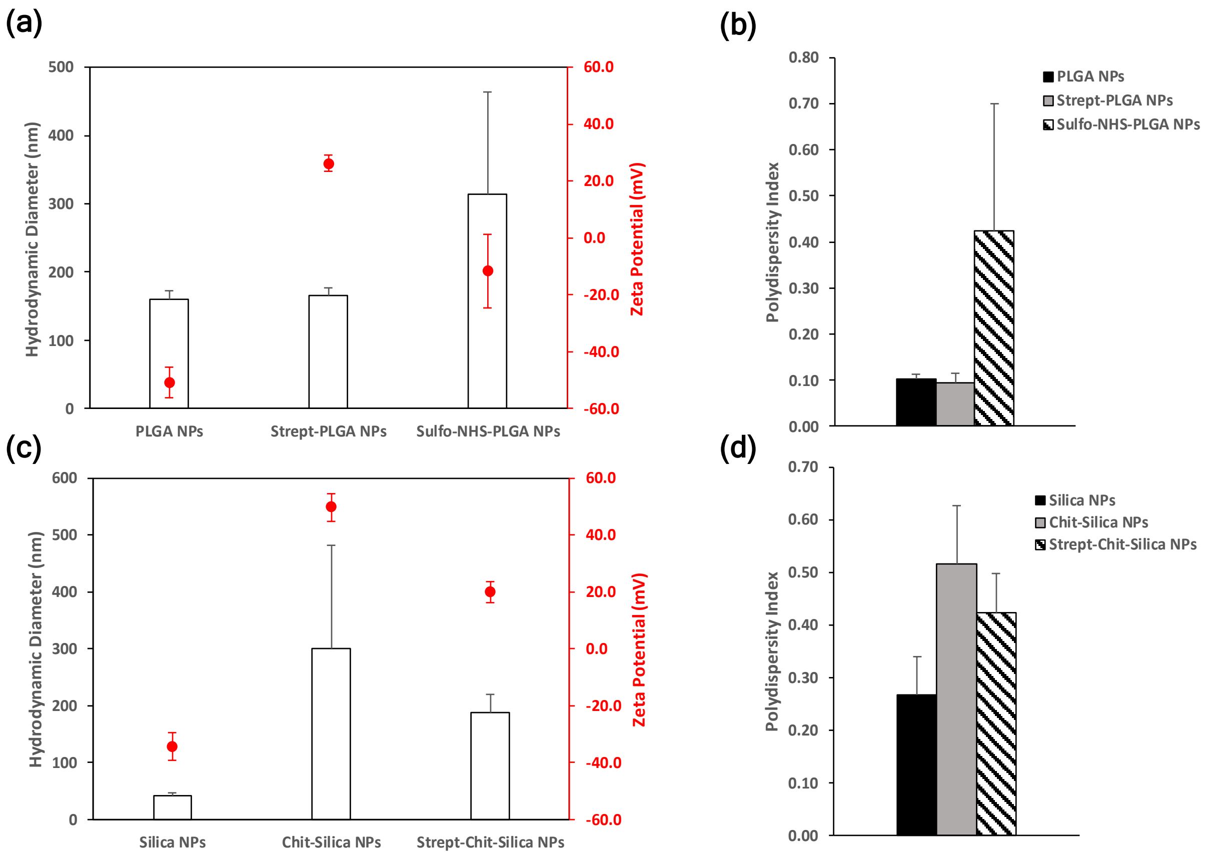


**Figure S3. Dynamic Light Scattering Measurements of Nanoparticle Suspensions Pre- and Post-Functionalization. (a-b)** Hydrodynamic diameter, zeta potential, and polydispersity index of PLGA nanoparticles (PLGA NPs), streptavidin-functionalized PLGA nanoparticles (Strept-PLGA NPs), and sulfo-NHS functionalized PLGA nanoparticles (Sulfo-NHS-PLGA NPs). **(c-d)** Hydrodynamic diameter, zeta potential, and polydispersity index of silica nanoparticles (Silica NPs), chitosan-coated silica nanoparticles (Chit-Silica NPs), and streptavidin-functionalized chitosan-coated silica nanoparticles (Strept-Chit-Silica NPs).
